# Genome-wide association study of appendicular lean mass in UK Biobank cohort

**DOI:** 10.1101/643536

**Authors:** Yu-Fang Pei, Yao-Zhong Liu, Xiao-Lin Yang, Hong Zhang, Gui-Juan Feng, Lei Zhang

**Affiliations:** Department of Epidemiology and Health Statistics, Medical College of Soochow University, Jiangsu, PR China; Jiangsu Key Laboratory of Preventive and Translational Medicine for Geriatric Diseases, School of Public Health, Medical College of Soochow University, Jiangsu, PR China; Department of Biostatistics and Data Science, Tulane University School of Public Health and Tropical Medicine, New Orleans, LA, USA; Center for Genetic Epidemiology and Genomics, School of Public Health, Medical College of Soochow University, Jiangsu, PR China

**Keywords:** appendicular lean mass, GWAS, UK Biobank

## Abstract

Lean body mass (LBM), an important physiological measure, has a strong genetic determination. To clarify its genetic basis, a large-scale genome-wide association study (GWAS) of appendicular lean mass (ALM) was conducted in 450,580 UK Biobank subjects. A total of 717 variants (p<5×10^−9^) from 561 loci were identified, which were replicated across genders (achieving p<5×10^−5^ in both genders). The identified variants explained ~11% phenotypic variance, accounting for one quarter of the total ~40% GWAS-attributable heritability. The identified variants were enriched in gene sets related to musculoskeletal and connective tissue development. Of interest are several genes, including *ADAMTS3*, *PAM*, *SMAD3* and *MEF2C*, that either contain multiple significant variants or serve as the hub genes of the associated gene sets. Polygenic score prediction based on the associated variants was able to distinguish subjects of high from low ALM. Overall, our results offered significant findings on the genetic basis of lean mass through an extraordinarily large sample GWAS. The findings are important to not only lean mass *per se* but also other complex diseases, such as type 2 diabetes and fracture, as our Mendelian randomization analysis showed that ALM is a protective factor for these two diseases.

## Introduction

Lean body mass (LBM) is an important physiological index. The decline of LBM with aging, also known as sarcopenia, is a critical factor for functional impairment and physical disability and a major modifiable cause of frailty in the elderly [1, 2]. LBM is associated with bone mineral density (BMD), and hence may be also relevant to risk for osteoporosis [3]. Other LBM-related conditions include dysmobility syndrome [4], sarcopenic obesity [5], and cachexia [6]. Overall, sarcopenia was responsible for an increased risk of mortality, with a hazard ratio of 1.29 to 2.39 [7].

LBM has a significant genetic component, as evidenced by a high heritability of 50% to 80% as observed in twin studies [8, 9]. However, findings on specific genes for human lean mass variation remain limited even with the powerful genome-wide association study (GWAS) approach. A key reason for the limited findings, as in other human complex traits, is the modest sample size used in most GWAS so far performed in LBM [10–14], resulting in few SNPs (single nucleotide polymorphisms) identified with genome-wide significance.

As a notable example, a recent large meta-analysis of GWAS amassed 20 cohorts of European ancestry with a total sample size of >38,000 for whole body lean mass (WBM) and of >28,000 for appendicular lean mass (ALM) [15]. However, despite of the large sample used, the percent variance explained by the identified SNPs was still only 0.23% and 0.16% for WBM and ALM, respectively, suggesting that most of the heritability of LBM was still undetected. Therefore, even with such a large GWAS meta-analysis, it is still necessary to boost the sample size further so as to enhance the statistical power for detecting more causal SNPs underlying LBM.

Here in this study, with a sample containing ~half-million subjects of European origin from UK Biobank (UKB), we performed a GWAS of appendicular lean mass (ALM). At the stringent genome-wide significance level (p<5×10^−9^), we identified >700 variants that were replicated across genders. Our findings revealed a large number of genetic variants for LBM and contributed to the characterization of the genetic architecture of this important complex trait. Through this GWAS we demonstrated the power for mapping the genetic landscape of common human complex traits/diseases using extraordinarily large samples.

## Results

Basic characteristics of the studied UKB sample are listed in **Supplementary Table 1**. In this study, we quantified appendicular lean mass (ALM) by appendicular fat-free mass measured by electronic impedance. This measurement of lean mass is reliable based on its strong correlation with ALM measured by DXA in 4,294 UKB subjects (with a Pearson’s correlation coefficient of 0.96, p<2.2×10^−16^).

### Main association results

Raw ALM was adjusted with appendicular fat mass (AFM) and the adjusted ALM (ALM_adj_) was the phenotype used for the GWAS. Following quality control (QC) of both ALM_adj_ and genome-wide genotypes, data from 19.4 million variants with minor allele frequency (MAF) >0.1% and imputation quality score >0.3 were available in 244,945 female and 205,635 male subjects.

In each gender group, additive effect of each variant was tested on ALM_adj_ with BOLT-LMM [16], controlling for age, age^2^, height and height^2^. The genomic inflation factor showed notable inflation in both gender groups (λ_female_=1.92, λ_male_=1.77). To examine observed inflation for potential polygenic effects and other biases, linkage disequilibrium score regression (LDSC) analysis was performed [17]. The estimated mean chi-square and intercept were 2.34 and 1.12 for females, and 2.53 and 1.15 for males, corresponding to an attenuation ratio (AR) of 0.098 and 0.090, respectively. The AR estimates are comparable to those estimated in the subset of 369,968 unrelated British white subjects (0.090 and 0.074 for females and males, respectively), who were extracted from the total sample.

Using BOLT-REML [18], GWAS-attributable heritability was estimated, which was 0.381 (s.e 3.30×10^−3^) and 0.394 (s.e 3.80×10^−3^) in females and males, respectively. LDSC estimated a genetic correlation coefficient as high as 0.90 (s.e 0.01) between the two genders, implying that most GWAS-attributable heritability was shared across genders.

Given the shared heritability across genders, across-gender meta-analysis was performed with the inverse variance weighted fixed-effects model to combine the gender-specific GWAS results. The meta-analysis signals have an AR of 0.115 (mean chi-square=3.69, intercept=1.31). Genome-wide significance (GWS) level was set to α=5×10^−9^, and a suggestive significance level was set to α=5×10^−5^. An association was declared to be “replicated” if it is 1) significant at the GWS level in the across-gender meta-analysis and 2) significant at the suggestive level within each gender.

Based on the above criteria, a total of 589 loci were identified at GWS level in across-gender meta-analysis (p <5×10^−9^), which were also replicated (p <5×10^−5^) across genders. To check potential linkage disequilibrium (LD) among these loci, LD analysis was performed on 589 lead variants (each from one of the loci). It was found that 47 lead variants are not in linkage equilibrium with each other (LD r^2^>0.1) due to long-range LD. After removing 28 loci, the lead variants in the remaining 561 loci were all in linkage equilibrium (LD r^2^ <0.1). Therefore, these 561 loci were treated as independent loci for downstream analysis.

Approximate conditional association analysis and across-gender meta-analysis were recursively performed, which further identified an additional set of 156 conditionally significant variants (p<5×10^−9^ in across gender meta-analysis) that were replicated across genders (p<5×10^−5^ within each gender). These additional variants were also in linkage equilibrium (LD r^2^<0.1) with the lead variants of the 561 loci.

In total, 717 (i.e., 561+156) independent variants from 561 distinct loci were associated with ALM_adj_ (**Supplementary Table 2**). Among the 717 lead variants, 172 achieved the strongest significance level (p<5×10^−9^) in both genders (categorized here as the Tier 1 variants). Also, 144 variants achieved p values <5×10^−9^ in females, and p values < 5×10^−5^ in males; 62 variants achieved p values < 5×10^−9^ in males, and p values < 5×10^−5^ in females (categorized here as the Tier 2 variants). At last, 339 variants achieved p values < 5×10^−5^ in both genders and p values < 5×10^−9^ in across-gender meta-analysis (categorized here as the Tier 3 variants).

Of the above identified loci, 17 were reported by GWAS or meta-analysis of DXA-derived lean mass [13, 15, 19]; 104 were reported by a study of electronic impedance measured lean mass in a subset UKB cohort subjects (N=155,961) [20].

We also evaluated the overlap of the identified loci with those identified for several obesity traits, including body mass index (BMI), waist circumference (WC), WC adjusted for BMI (WC_adj_BMI), waist-hip ratio (WHR) and WHR adjusted for BMI (WHR_adj_BMI). SNPs in 302 loci (defined as the lead SNP + 500 kb flanking at each side) showed association with one or more obesity traits at the conventional significance level 5.0×10^−8^, while no trait was associated with the remaining 259 loci, demonstrating their novelty and possibly, specificity to lean, but not fat mass.

### Gender heterogeneity/specificity

In addition to the 561 loci that are replicated across genders, our analysis also identified 152 loci that are significant (p<5×10^−9^) in across-gender meta-analysis but not significant at the suggestive level (p<5×10^−5^) in either gender group (**Supplementary Table 3**). These loci may represent gender specific signals pending further replication.

Of the 717 identified variants (of the 561 loci), 109 (15.2%) have a high level across-gender meta-analysis heterogeneity (I^2^>50%), all (except one) of which belong to Tiers 1 or 2 variants. A statistical test on gender difference in allele effect size showed that the difference is significant in only 2 SNPs, rs2972156 (p_diff_=2.49×10^−12^) and rs1933801 (p_diff_=4.65×10^−6^), after accounting for multiple testing (α=0.05/717=6.97×10^−5^), suggesting that almost all of the identified variants may have similar effect sizes across genders. The two SNPs (rs2972156 and rs1933801) with different effect size between genders achieved p values of 1.30×10^−46^ and 2.40×10^−26^, respectively, in males and p values of 4.20×10^−7^ and 1.30×10^−6^, respectively, in females.

### Heritability distribution

The 717 identified variants include 654 common variants (MAF>5%), 52 less common variants (5%≥MAF>1%) and 11 rare variants (MAF≤1%). Collectively, these variants explain 10.82% phenotypic variance in the total sample, most of which (9.91%) is accounted for by common variants. As expected, variants with a smaller MAF generally have a larger per allele effect size **(Figure 1**). For example, the average per allele effect size in rare variants (mean 0.11, s.d 0.05) is 6-fold larger than common variants (mean 0.02, s.d 0.007).

**Figure 1.**
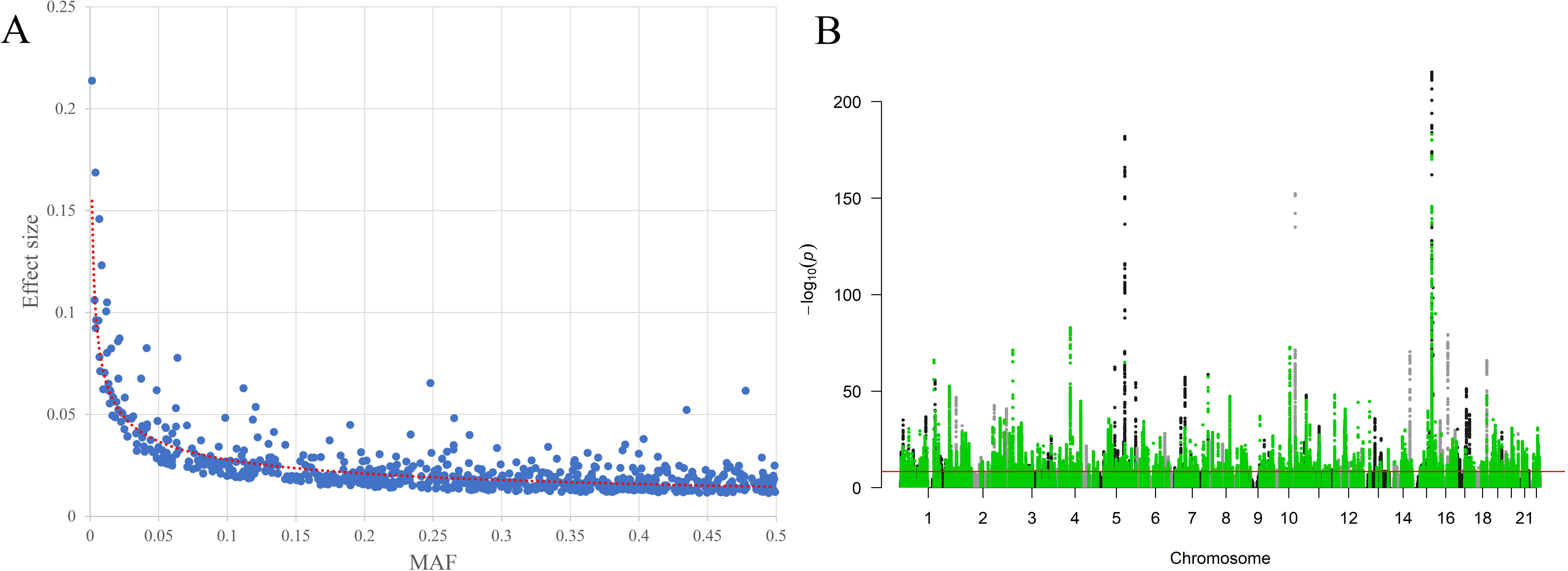
Main association results. **Figure 1A**, Per allele effect size versus minor allele frequency (MAF). X-axis is MAF at the 717 identified variants and y-axis is per allele effect size (regression coefficient). **Figure 1B**, the Manhattan plot of the meta-analysis combining both genders. The horizontal red line indicates the genome-wide significance level (alpha= 5×10^−9^) in −log_10_ scale. All novel loci were marked in green.

Applying the stratified LDSC analysis, the explained heritability was partitioned into 24 functional categories [21]. Statistically significant enrichments were observed for 19 functional categories (p<0.05/24, **Figure 2**). In line with the observations by Finucane et al. [21], regions conserved in mammals showed the strongest enrichment of any category, with 2.6% of SNPs explaining an estimated 34.5% of SNP heritability (enrichment ratio (EA)=13.2, P=3.39×10^−19^). Other categories with significant enrichment included coding regions (EA=8.8, P=1.76×10^−7^), 3’UTR (EA=5.7, P=3.73×10^−4^), transcription starting site (EA=5.1, P=1.71×10^−5^), and H3K9ac histone marks (EA=5.1, P=2.07×10^−15^). Neither promoter nor 5’-UTR region showed significant enrichment, though 5’-UTR region had a high estimate of EA (15.5, p=0.03).

**Figure 2.**
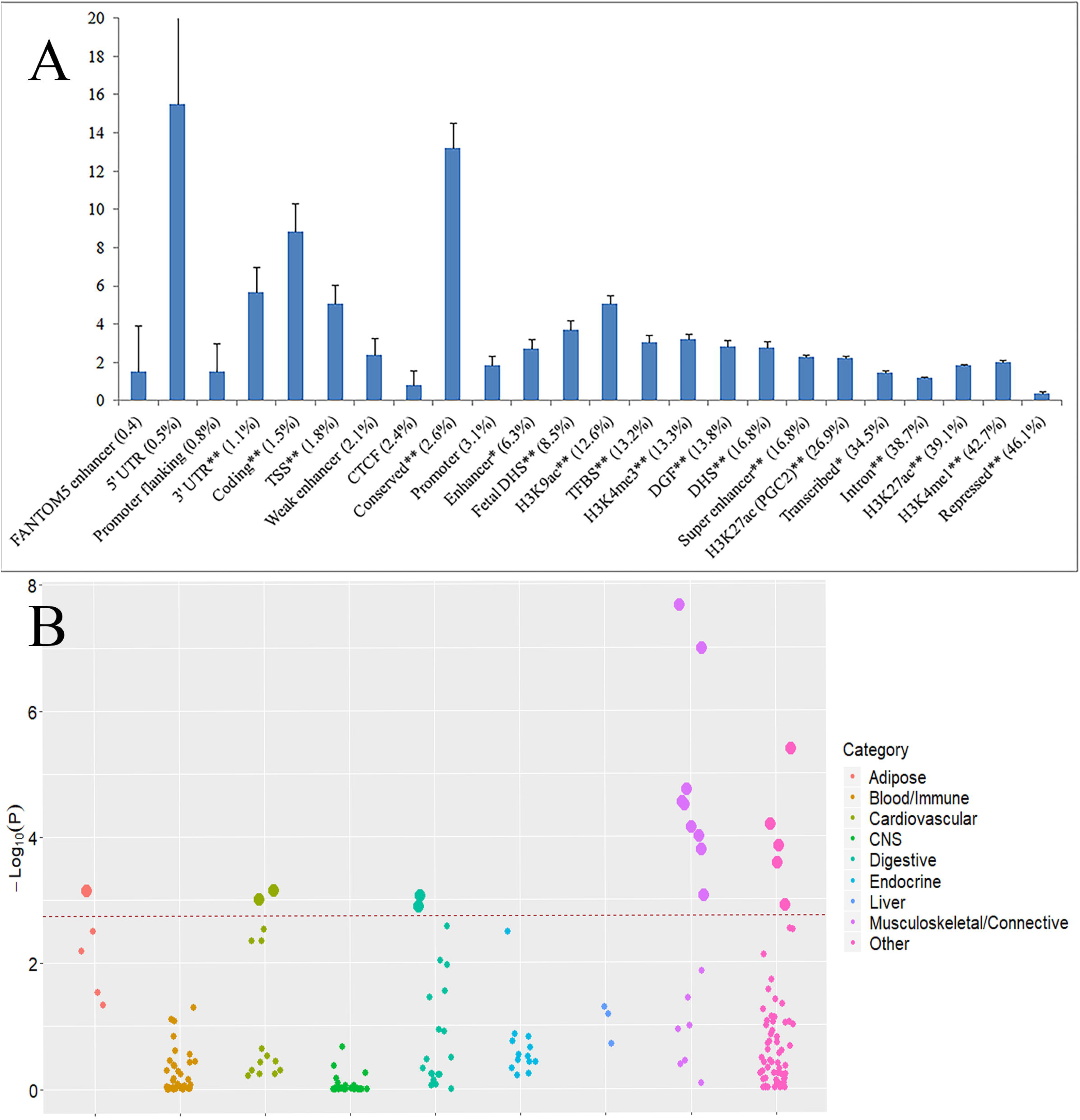
Heritability enrichment in different functional annotations and tissues. **Figure 2A** is enrichment of genome-wide association signals in 24 main annotations using LDSC regression. Y-axis represents the ratio of phenotypic variance explained by variants in a particular annotation category against that explained in the remaining regions. Error bars represent jackknife standard errors around the estimates of enrichment. A single asterisk indicates significance at p<0.05 after Bonferroni correction for the 24 hypotheses tested, and two asterisks indicates significance at p<0.01. **Figure 2B** is enrichment of genome-wide association signals in 206 cells/tissues from two different databases (Franke lab dataset and GTEx consortium dataset). The total cells/tissues were divided into 9 categories. Each dot represents a specific cell/tissue and the tissues passing the cutoff of FDR < 5% at −log10 (p) = 2.75 were marked in large.

A new function of the stratified LDSC method was used to assess focal tissues for heritability enrichment [22], using two gene expression datasets [23, 24]. A total of 19 tissues/cells are enriched at a false discovery rate (FDR) <5% (**Figure 2**). About half (9) of them belong to musculoskeletal and connective system, including cartilage, chondrocytes, osteoblasts, fibroblasts, smooth muscle, myometrium, cervical vertebrae, synovial membrane and stromal cells.

### Candidate genes prioritization

To prioritize candidate genes at the associated loci, we used multiple analytical strategies. A set of credible risk variants (CRVs) at each locus were defined as variants with high LD with the lead variant (r^2^>0.8). A total of 17,968 CRVs were defined (**Supplementary Table 4**). Based on the CRVs, 6 types of supporting evidence were used to prioritize 1,337 candidate genes. (**Supplementary Tables 5-10**).

A number of genes have multiple lines of supporting evidence. Peptidylglycine Alpha-Amidating Monooxygenase (*PAM*) at 5q21.1, in particular, has all lines of supporting evidence. This locus contains two independent signals. The first is a mis-sense rare SNP rs78408340 (MAF=0.01%, p=6.10×10^−10^) inside PAM, and the second is a common SNP rs400596 located between *PAM* (129.5 kb from *PAM*) and *SLC06A1* genes (237.2 kb from *SLC06A1*). Polymorphisms at rs400596 are associated with the *PAM* expression level in whole blood (p=2.51×10^−21^) [23] and associated with its protein level in peripheral blood (p=p=2.51×10^−30^) [25]. *PAM* is also prioritized by both SMR [26] and DEPICT [27], strengthening its functional relevance.

### Comparison between imputation and sequencing-based association signal

Of the 717 identified variants, 42 are mis-sense coding ones. Forty of them, including 7 rare ones, are available in the recently released UKB exome sequencing data that contain a subset of ~50,000 subjects from the whole UKB cohort. Using a set of 45,554 unrelated European subjects who were both genotyped/imputed and sequenced, we compared the imputation-based association results with exome sequencing based results. The 7 rare variants appeared to have limited imputed dosage variation hence their imputation association p-values were not able to derive. In the sequencing data, 3 of these 7 variants were nominally significant (p<0.05, **Supplementary Table 11**), suggesting limited power in imputation-based association analysis (compared with sequence-based analysis) for rare variants. This limited power may be alleviated by increased sample size since in the whole UKB cohort these 7 rare variants achieved significant p values in imputation-based association analysis.

Of the remaining 33 variants, the imputation-based and sequencing-based p-values were highly concordant. For example, the imputation-based p-values are within 2-fold difference of the sequencing-based p-values for up to 29 variants. Overall, these observations support that imputation-based association signals are close to the real sequencing-based association signals in a large sample. Therefore, imputation based GWAS may be able to identify true associations, even for those of rare variants.

### Mis-sense variants and the associated genes

As mentioned above, of the 717 identified variants, 42 are mis-sense coding ones. Majority of these 42 mis-sense mutations are predicted to be deleterious according to more than one bioinformatics tool including PolyPhen2 [28], SIFT [29], PROVEAN [30] and Fathmm [31] (**Supplementary Table 12**), supporting their functional relevance.

Mis-sense mutations are enriched among rare variants. Eight of the 11 rare variants are mis-sense mutations, which is in clear contrast to 34 mis-sense mutations among the remaining 706 variants (odds ratio (OR)=55.37, Fisher’s exact test p=7.11×10^−9^). Evidence of the enrichment became stronger by comparing 21 mis-sense mutations from 63 rare or less common variants vs. 22 mis-sense mutations from 654 common variants (OR=14.36, Fisher’s exact test p=5.75×10^−13^), suggesting that low frequency mutations are more likely to play a direct role in changing protein function.

Genes containing mis-sense variants are listed in **Table 1**. In particular, the *ADAMTS3* gene contains 3 rare or less common mis-sense variants (rs141374503 MAF=0.4% p=2.02×10^−27^; rs150270324 MAF=1.3%, p=2.36×10^−14^, and rs139921635 MAF=2.4%, p=4.06×10^−15^). In addition, it also contains multiple non-mis-sense variants, including 3 conditionally significant variants in its intron region: rs72653979 (MAF=7.8%, p=9.51×10^−11^), rs78862524 (MAF=5.5%, p =3.24×10^−23^) and rs769821342 (MAF=3.2%, p =1.27×10^−14^), and 2 in its flanking inter-genic region: chr4:73496010 (MAF=47%, p=3.87×10^−21^) and rs10518106 (MAF=6%, p=1.16×10^−83^). Though these SNPs are 367.0 kb apart at most, they are in linkage equilibrium with each other (LD r^2^<0.1). Together, the 8 variants from the *ADAMTS3* gene explain 0.18% of phenotypic variance, making this region the most contributive locus.

**Table 1.**
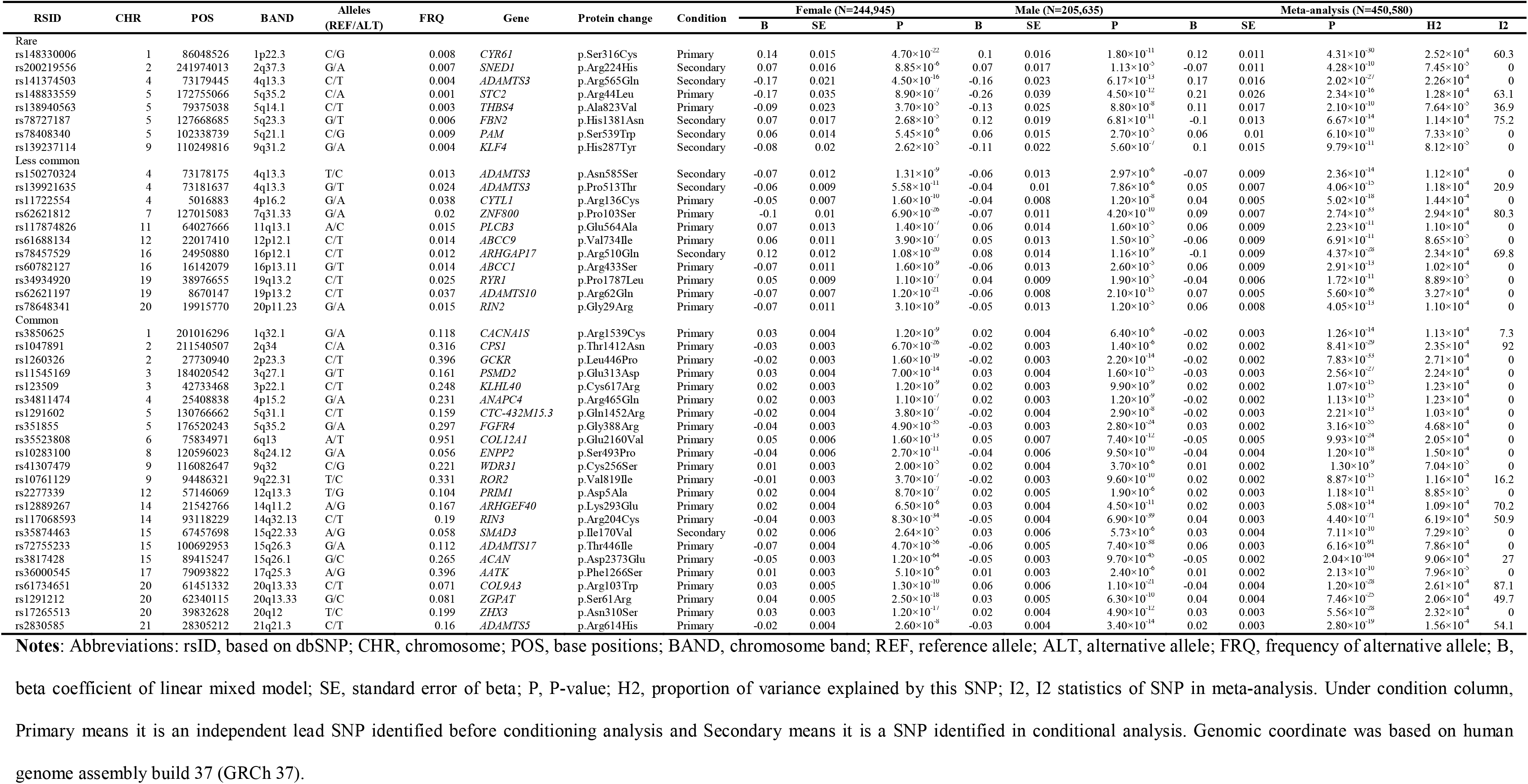
Association results of 42 mis-sense variants.

### Gene-based and gene set enrichment analyses

A total of 3,101 genes were significant at the gene-based genome-wide significance level (α=0.05/19,098=2.62×10^−6^, **Supplementary Table 13**), and 85 gene sets were significant at the gene set significance level (α=0.05/10,655=4.69×10^−6^, **Supplementary Table 14**).

The most significant gene set is GO:0001501 ‘skeletal system development’ (p=8.88×10^−24^), followed by GO:0031012 ‘proteinaceous extracellular matrix’ (p=5.01×10^−14^), GO:0061448 ‘connective tissue development’ (p=1.10×10^−13^), GO:0048705 ‘skeletal system morphogenesis’ (p=9.33×10^−13^) and GO:0031012 ‘extracellular matrix’ (p=1.05×10^−12^). Additional gene sets with known function related to musculoskeletal and connective system, such as GO:0051216: ‘cartilage development’ (p=3.30×10^−12^) and GO:0042692 ‘muscle cell differentiation’ (p=1.45×10^−6^), were also identified.

Genes involved in multiple gene sets are likely to act as hub genes and may play a central regulatory role. From the list of significant gene sets, the most frequently involved gene is *SMAD3* (gene-based association p=8.11×10^−42^), which was involved in 46 out of the 85 significant gene sets. It was followed by *SOX9* (p=0.05, in 44 gene sets), *MEF2C* (p=2.83×10^−9^, in 42 gene sets) and *BMP4* (p=0.15, in 42 genes). All these 4 genes were reported by previous studies as important candidate genes for muscle development [32–35]. However, *SOX9* is only nominally significant and *BMP4* is not significant at single gene level, indicating that the significant pathway signals may not be contributed by the two genes. Altogether, there are 34 genes, each of which was involved in more than 30 of the 85 significant gene sets.

Protein-protein interaction (PPI) analysis using these 34 hub genes connects them into a tight interactional network (**Figure 3**). This network contains multiple genes that are important for skeletal muscle development, such as TGF signaling pathway genes (*TGFB1*, *TGFB2* and *TGFBR2*), BMP signaling pathway genes (*BMP2* and *BMP4*) and SMAD family genes (*SMAD1*, *SMAD2*, *SMAD3* and *SMAD4*).

**Figure 3.**
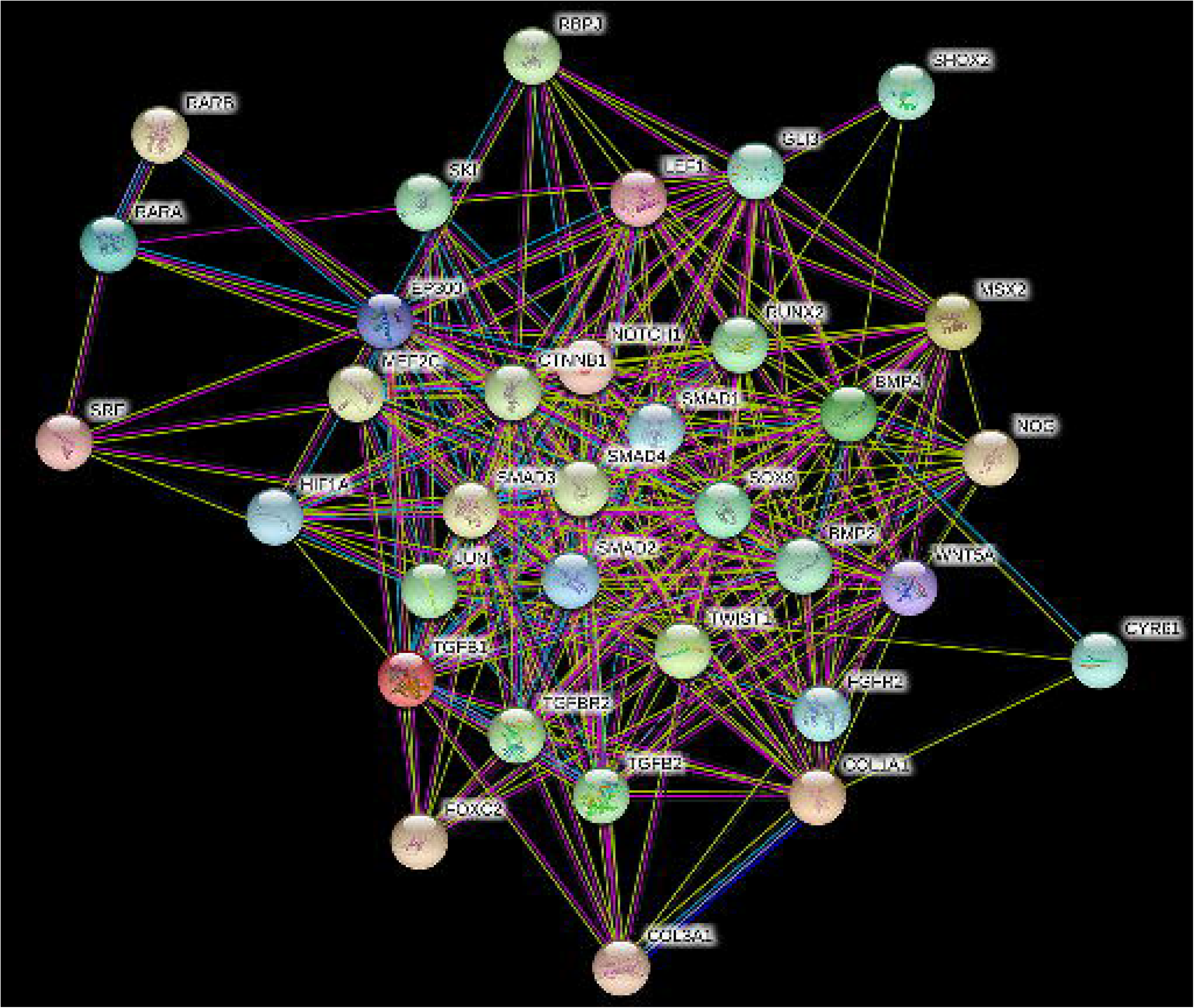
Protein-protein interactional network. Thirty-four genes over-represented in 85 significant pathways were selected to construct a protein-protein interaction network with STRING, which bases the construction on knowledge of gene co-expression, text-mining, and others.

### Polygenic risk score profiling

To assess the ability of the GWAS findings to predict ALM, a polygenic scoring analysis was performed in the subset of 369,968 unrelated British white subjects from the UKB cohort. Three quarters of the subjects (277,762 participants, including 149,329 females) were randomly selected as the training sample, with the remaining subjects (92,206 participants, including 49,660 females) as the validation sample.

The training sample identifies 72,456 variants that achieved a p-value <1×10^−5^ for association with ALM_adj_. Using these variants as predictor, the predicted genome-wide polygenic score (GPS) and the real phenotype residual in the validation sample are significantly correlated (Pearson’s correlation coefficient 0.22, 95% CI (0.21, 0.22), p<2.2×10^−16^). Mean phenotype residuals in the top tail are significantly higher than that in the bottom tail of the GPS distribution (**Figure 4**). For example, the predicted top 1% subjects have an increased average residual of 1.16 than the predicted bottom 1% participants (0.57 (s.d 0.96) vs. −0.59 (s.d 0.94)), corresponding to an 1.69 kilo-gram (kg) increase of raw ALM (24.61 kg (s.d 5.89 kg) vs. 22.92 kg (s.d 5.27 kg)). In the female group, the predicted top 1% participants have on average 1.39 kg increase of raw ALM than the predicted bottom 1% participants (20.26 kg (s.d 2.75 kg) vs. 18.87 kg (s.d 2.45 kg)). In males, the increase is 2.29 kg (29.82 kg (s.d 4.18 kg) vs. 27.53 kg (s.d 3.56 kg)). These results demonstrate that the GPS prediction based on the current GWAS finding is capable of identifying subjects of high or low levels of ALM.

**Figure 4.**
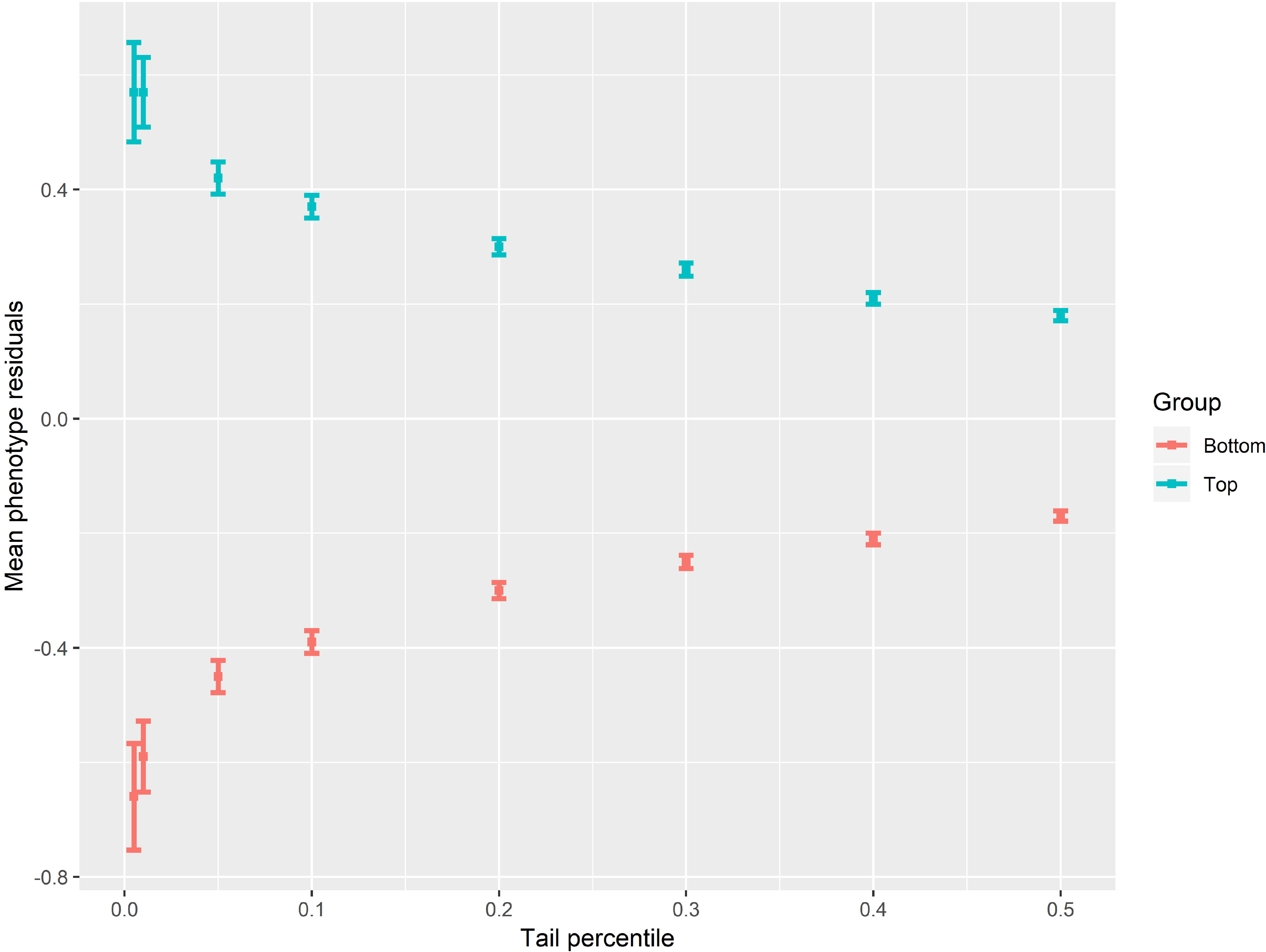
Polygenic score prediction. A total of 277,762 subjects were randomly selected as the training sample, and another independent 92,206 subjects were selected as the validation sample. The variants achieving a p-value of <1×10^−5^ in the training sample were selected and used for prediction in the validation sample via LDpred approach. Subjects in the two extreme tails of the predicted genome-wide polygenic score (GPS) distribution were compared in terms of raw phenotype mean (after correction). X-axis represents the fraction of subjects drawn from both extreme tails of the predicted GPS distribution. Y-axis represents mean ALM_adj_ (±95% confidence interval).

### Genetic correlations with other traits

To test whether lean mass has a shared genetic etiology with other diseases and relevant traits, a genetic correlation analysis was performed with the LDSC method [17]. Here, ALM studied in our study is strongly genetically correlated with DXA-derived whole body lean mass and the ALM, which were studied by a previous GWAS meta-analysis [15] (r_g_=0.87 and 0.78) (**Figure 5**). Furthermore, ALM is modestly correlated with BMI (r_g_=0.31). However, the correlation with BMD is low (r_g_=0.05). ALM is most negatively correlated with BMI adjusted leptin (r_g_=−0.41). It is also negatively correlated with body fat (r_g_=−0.17), suggesting a reverse developmental direction towards lean and fat mass.

**Figure 5.**
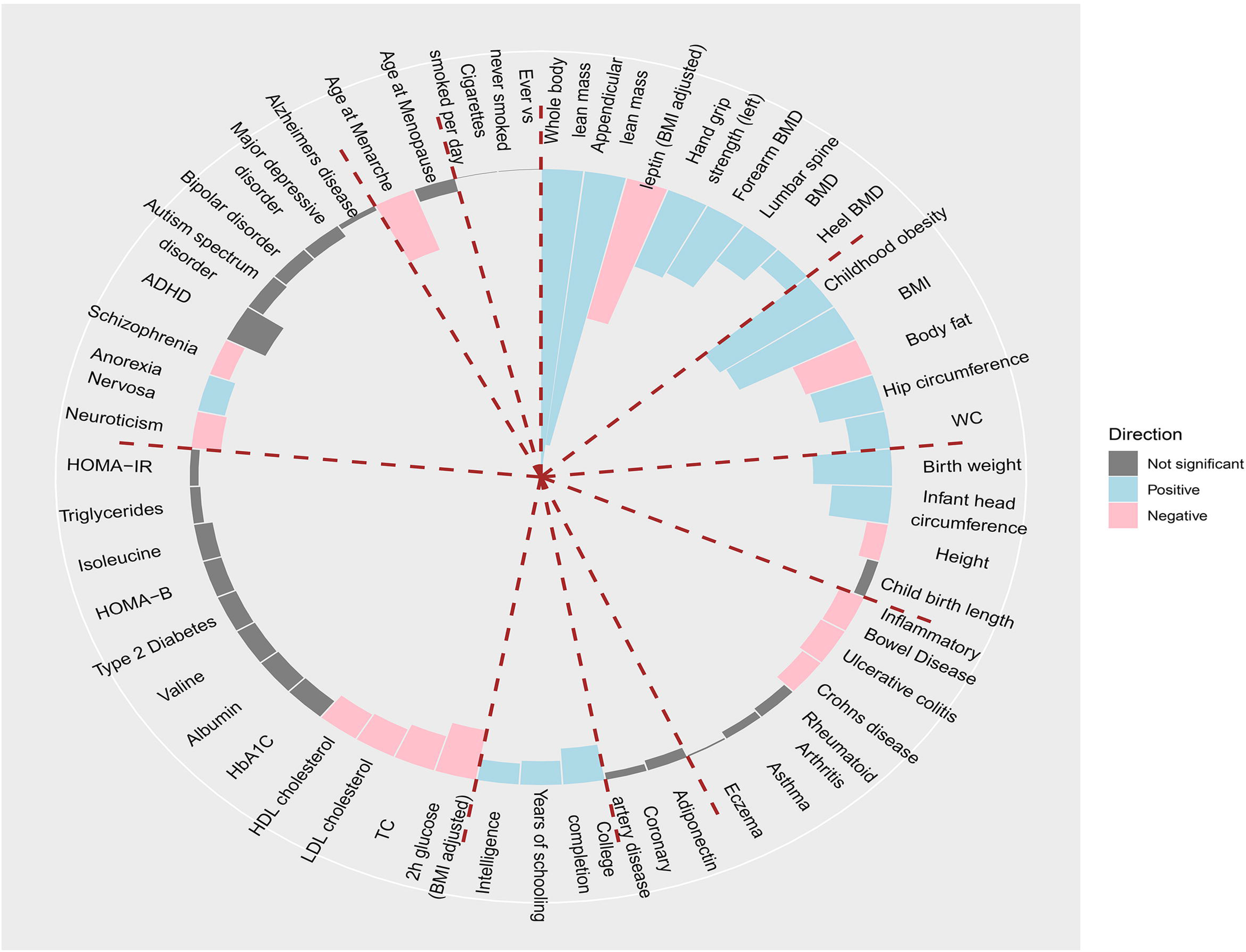
Genetic overlap with other traits. Genetic correlations (r_g_) between ALM_adj_ and 51 traits and diseases were estimated. LD Score regression tested genome-wide SNP associations for these participants against similar data for various other traits and diseases containing Musculoskeletal system, anthropometrics, obesity, cognition, metabolism, psychiatry, reproduction and neuropsychiatric outcomes. Error bars represent standard errors on these estimates. Blue bars represent significantly positive correlation at the nominal level p<0.05; pink bars represent significantly negative correlation (p<0.05); grey bars represent non-significant correlation.

### Mendelian randomization analysis

To investigate whether ALM is causally linked with other complex diseases, a Mendelian randomization analysis was performed with GSMR [26]. Ten diseases from a variety of categories were chose for evaluation, including fracture [36], type 2 diabetes (T2D) [37], asthma [38], insomnia [39], inflammatory bowel disease (IBD) [40], smoking addiction [41], coronary artery disease (CAD) [42], amyotrophic lateral sclerosis (ALS) [43], bipolar disorder [44] and autistic spectrum disorder (ASD) [45]. At the corrected significance level 5×10^−3^ (0.05/10), ALM is causally associated with type 2 diabetes (T2D, p=4.38×10^−8^) and fracture (p=1.18×10^−3^), but not with any other disease (**Supplementary Table 15**). Specifically, a negative association is observed between ALM and both diseases, indicating that ALM is a protective factor for both diseases. For T2D, an increase in the unit of one s.d of ALM residual corresponds to a decreased OR of 0.91 (95% CI [0.88, 0.94]). For fracture, an increase in the unit of one s.d. of ALM residual corresponds to a decreased OR of 0.95 (95% CI [0.92, 0.98]).

## Discussion

The incapacity in GWAS to detect and replicate specific genetic variants for human complex traits, contradicting to a trait’s established high heritability, e.g., height, was formally recognized as the “missing heritability” problem a decade ago [46, 47]. An explanation is the so called “polygenic model”, where hundreds or even thousands of common SNP variants act additively, with each contributing only a “tiny” fraction of the trait variation. The effect of each individual variant is so small that a GWAS with a limited sample size (n<20,000) may be extremely difficult, if not impossible, to detect (let alone replicate) a variant at the genome-wide significance threshold.

The polygenic model was supported by the genome-wide complex trait analysis (GCTA), where trait similarity among unrelated subjects was correlated with and explained to a large fraction by similarity of common SNPs at genome-wide scale [48]. Furthermore, with sample sizes at the scale of hundreds of thousands, two GWAS indeed identified at genome-wide significance ~700 variants for adult height [49] and >100 loci for schizophrenia [50]. The successful stories offer a promising prospect for a GWAS with an extraordinarily large sample size to ultimately unravel the puzzling genetic architecture for human complex traits and common diseases.

In this study of lean mass with around half million subjects, the largest sample used for GWAS of lean mass so far, a successful endeavor was accomplished again. More than 700 variants were identified at the significance of genome-wide scale (p<5×10^−9^). In particular, more than half of these variants achieved genome-wide significance (p<5×10^−9^) in one gender and were replicated also in another gender (p<5×10^−5^). Overall, these >700 variants contributed ~11% of ALM variation, again, the largest explainable fraction of variation for lean mass reported so far in a GWAS.

Our findings of >700 variants are expected for a complex trait with a high heritability, particularly considering another trait with comparable heritability, height, which detected also ~700 variants [49]. Interestingly, the majority loci in previous smaller GWAS [13] or meta-analysis [15] of lean mass are also significant in the present study, providing replication evidence from independent samples.

The functional relevance of our identified variants is supported by the gene set enrichment analysis, where GO terms, including GO:0001501 ‘skeletal system development’, GO:0061448 ‘connective tissue development’, GO:0051216 ‘cartilage development’ and GO:0042692 ‘muscle cell differentiation’, are among the top gene sets of significance. Specifically, the “hub genes” involved in these terms are tightly connected into a network that contains TGF pathway genes, BMP pathway genes and SMAD family genes, which are all important musculoskeletal development genes/pathways. This finding is concordant with developmental biology since cells from bone, cartilage, muscle and fat share the same progenitor, the mesenchymal stem cells, and pleiotropy of muscle and bone is well recognized in both humans [51] and animal models [52].

Among the variants identified, those of several genes, such as *SMAD3*, *MEF2C*, *ADAMTS3* and *PAM*, are interesting and may need further investigation. The first two genes are the hub genes involved in half of the significant enriched gene sets. The third gene, *ADAMTS3*, contains 8 variants, including 3 rare or less common mis-sense mutations, which in total explains ~0.2% of ALM variation. The fourth gene, *PAM*, has multiple lines of supporting evidence for its regulatory roles, e.g., containing a mis-sense rare SNP rs78408340 (MAF=0.01%, p=6.10×10^−10^). An intergenic variant, rs400596, is associated with the PAM expression level in whole blood (p=2.51×10^−21^) [23] and associated with its protein level in peripheral blood tissue (p= p=2.51×10^−30^) [25]. These genes may represent the next candidates for functional and mechanistic analysis of lean mass regulation.

In summary, we performed a GWAS using ~half-million subjects for lean mass. Owing to its high statistical power, our study identified a large number of variants mapped to GO terms with functional relevance to musculoskeletal development. The explained variation of ~11% of lean mass by the identified variants represents a significant leap in revealing the “hidden” heritability of this complex trait using GWAS. Our findings’ translational value is marked by lean mass’ importance to other complex diseases, such as type 2 diabetes and fracture, as our Mendelian randomization analysis showed that ALM is a protective factor for these two diseases. Overall, our study provides another example, where GWAS of substantially increased sample size may lead a way to ultimately and thoroughly delineate genetic architecture of human complex traits. This epitomizes the value of big data in genetic research of humans.

## Materials and Methods

### Study participants

Study sample came from the UK Biobank (UKB) cohort, which is a large prospective cohort study of ~500,000 participants from across the United Kingdom, aged between 40-69 at recruitment. Ethics approval for the UKB study was obtained from the North West Centre for Research Ethics Committee (11/NW/0382), and informed consent was obtained from all participants. This study (UKB project #41542) was covered by the general ethical approval for the UKB study.

All the included subjects are those who self-reported as white (data field 21000). Subjects who had a self-reported gender inconsistent with the genetic gender, who were genotyped but not imputed or who withdraw their consents were removed. The final sample consisted of 450,580 subjects, including 244,945 females and 205,635 males.

### Phenotype and modeling

Body composition was measured by bioelectrical impedance approach. Appendicular lean mass (ALM) was quantified by the sum of fat-free mass at arms (data fields 23121 and 23125) and at legs (data fields 23113 and 23117). Appendicular fat mass (AFM) was quantified by the sum of fat mass at arms (data fields 23120 and 23124) and at legs (data fields 23112 and 23116). In each gender, covariates including AFM, age, age^2^, height and height^2^ were tested for significance in association with ALM using a step-wise linear regression model implemented in the R function stepAIC. Raw ALM values were adjusted by the significant covariates, and the residuals were normalized into inverse quantiles of standard normal distribution, which were used for subsequent association analysis.

A small subset of 4,294 subjects also received a dual-energy X-ray absorptiometry (DXA) body composition scan, and hence their DXA-derived ALM is also available. Therefore, raw ALM derived from DXA and from electric impedance was compared in these subjects by Pearson’s correlation coefficient.

### Genotype quality control

Genome-wide genotypes for all subjects were available at 784,256 genotyped autosome markers, and were imputed into UK10K haplotype, 1000 Genomes project phase 3 and Haplotype Reference Consortium (HRC) reference panels. A total of ~92 million variants were generated by imputation. We excluded variants with MAF<0.1% and with imputation r^2^<0.3. As a result, ~19.4 million well imputed variants were retained for subsequent genetic association analysis.

### Genetic association analysis

In each gender group, we used BOLT-LMM to perform linear mixed model (LMM) analysis [16]. As the LMM analysis can adjust for population structure and relatedness, we included all eligible subjects into analysis, as recommended by BOLT [53]. We did not include principal components (PCs) of ancestry as covariates in the LMM analysis.

After sex-specific associations were analyzed, we meta-analyzed the summary statistics of the two genders by inverse-variance weighted fixed-effects model with METAL [54]. The genome-wide significance (GWS) level was set at α=5×10^−9^, to account for both common and rare variants. The variants that passed this threshold in across-gender meta-analysis were then checked for replicability across genders based on a suggestive significance level 5×10^−5^ in each gender. The suggestive level was set so as to account for multiple testing of presumed maximal number of 1000 independent loci (0.05/1000). An association was defined as “replicated” if the signal was significant at the GWS level (p<5×10^−9^) in the meta-analysis and was significant at the suggestive level (p<5×10^−5^) in both genders.

This declaration of a replicated association was approximately same as a two-stage design, where the first stage involves selecting variants at the suggestive level (p<5×10^−5^) in one gender and the second stage involves replicating the selected variants at the same significance level (p<5×10^−5^) in another gender. An association locus was defined as a genomic region of 500 kb to both sides of a significant lead signal.

Difference in effect size between female and male was examined by a two-tailed p-value from the z-score in the following equation

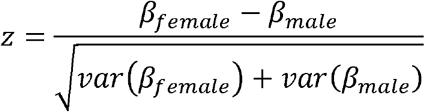

where *β*_*female*_ and *β*_*male*_ are regression coefficients for females and males, and var(·) are their variances, respectively.

### Conditional association analysis

To identify additional signals in regions of association, approximate joint and conditional association analysis was performed in each region using the GCTA tool [55].

From the UKB sample, a reference sample of 100,000 unrelated subjects was generated for estimating LD pattern for subsequent analyses. The unrelated subjects were inferred with KING software [56], from whom the 100,000 subjects of the reference sample were randomly drawn. Quality control (QC) procedures applied to the reference sample included Hardy-Weinberg equilibrium (p>1×10^−6^) and MAF>0.1%.

A recursive conditional association analysis was performed. In each iteration, an approximate conditional analysis conditioning on the current list of lead variants was performed in each gender, followed by an across-gender meta-analysis to combine the gender-specific results. Again, a significant replicated association was defined as achieving both a conditional meta-analysis GWS signal (p<5×10^−9^) and a conditional suggestive signal (p<5×10^−5^) in both genders. In addition, each such identified variant is required to be independent of all variants in the lead SNP list (LD r^2^<0.1). The variant with the lowest p-value among such identified ones was added into the list of lead variants. Iterations of the conditional analysis were run until no significant signal can be identified.

### Overlap with loci in previous GWAS of obesity traits

GWAS summary statistics for 5 obesity traits, including body mass index (BMI) [57], waist circumference (WC), WC adjusted for BMI (WC_adj_BMI), waist-hip ratio (WHR) and WHR adjusted for BMI (WHR_adj_BMI) [58], were downloaded from the GIANT consortium website. For each trait, SNPs located within all the 561 identified loci (lead SNP +500 kb flanking region at each side) were extracted from the GWAS summary statistics. Significance level for the obesity traits were set at the conventional level of 5.0×10^−8^.

### Exome sequencing association analysis

During the preparation of this manuscript, the UKB released exome-sequencing data on a selected subset of ~50,000 participants. We compared the exome-sequencing based association results with that based on genotype imputation. To accomplish this, we generated an unrelated sample consisting of subjects who were both exome-sequenced and genotype-imputed.

As the QC procedure, we removed subjects who were not self-reported as white, whose self-reported genders were inconsistent with their genetic genders, and who withdrew their consents. The KING software was used to select unrelated subjects based on pairwise kinship matrix for up to 2^nd^ degree relatedness [56]. The final sample consisted of 45,554 participants, including 24,740 females and 20,814 males.

Sequence variant coordinates, which were annotated to the GRCH38 assembly, were converted back to the GRCH37 assembly with Liftover (http://genome.ucsc.edu/cgi-bin/hgLiftOver). For each subject, variants that were missing in the sequenced data were set to missing in the imputed data as well. In both datasets, genetic association with normalized phenotype residuals was analyzed with PLINK2 [59]. The top 10 PCs were included as covariates to account for potential population stratification.

### Genetic architecture

BOLT-REML was used to estimate heritability tagged by all the analyzed variants [18]. LD score regression (LDSC) method was used to estimate the amount of genomic inflation due to confounding factors such as population stratification and cryptic relatedness [17]. Pre-computed LD scores from the 1000 Genomes project European subjects were used for estimation. The relative contribution of confounding factors was measured by attenuation ratio (AR), which is defined as (intercept-1)/(mean chi^2−1), where intercept and mean chi^2 are estimates of confounding and the overall association inflation, respectively [17].

To compare AR with that estimated on unrelated subjects, a maximal subset of unrelated subjects from the total sample being analyzed were generated. Specifically, KING was used to extract a subset of unrelated subjects [56]. The resulting unrelated sample included 369,968 participants (198,989 females and 170,979 males). In each gender, PLINK2 was used to perform genetic association analysis [59]. To account for genetic confounding, the top 10 PCs inferred from UKB were used as the additional covariates.

To calculate the variance explained by all independent lead variants, individual variant effect size was estimated with the formula 2*f*(1−*f*)β^2^, where *f* is allele frequency and β is regression coefficient associated with the variant.

### Enrichment analysis

Stratified LDSC was used to partition heritability from GWAS summary statistics into different functional categories [21]. The analysis was based on the ‘full baseline model’ created by Finucane et al. [21] from 24 publicly available main annotations that are not specific to any cell type. Significance level of enrichment was set at p < 2.08×10^−3^ (0.05/24).

The stratified LDSC was used to also assess the enrichment of heritability into specific tissues and cell types [22]. This method analyzes gene expression data together with GWAS summary statistics, for which, the two pre-compiled gene expression datasets in LDSC were used. The first one is the GTEx project v6p [23] and the second one is the Franke lab dataset [24]. The GTEx dataset contains 53 tissues with an average of 161 samples per tissue. The Franke lab dataset is an aggregation of publicly available microarray gene expression datasets comprising 37,427 human samples from 152 tissues. The total 205 (=53+152) tissues are classified into nine categories for visualization. Significance was declared at a false discovery rate (FDR)<5%.

### Candidate gene prioritization

In each associated locus, a set of credible risk variants (CRVs) were defined as those variants in strong LD with the lead variant (r^2^>0.8, including lead variant). LD r^2^ measure was estimated based on the 100,000 unrelated reference sample with LDstore [60]. Six sources of information was used to evaluate a gene's causality: 1) being nearest to the lead CRV; 2) containing a mis-sense coding CRV; 3) being a target gene for a cis-eQTL CRV; 4) being a target gene for a cis- protein QTL (cis-pQTL) CRV; 5) being prioritized by DEPICT analysis [27] and 6) being prioritized by SMR analysis [61].

Cis-eQTLs revealed by the GTEx (v7) project were accessed from the GTEx web portal (http://www.gtexportal.org/) [23]. Cis-eQTL information is available for over 50 tissues. We selected skeletal muscle and whole blood for our analysis. Cis-eQTL was searched within 500 kb distance from a target gene. Significant cis-eQTL was declared at p<5×10^−5^.

Cis-pQTL information was accessed from Sun et al. [25]. GWAS summary statistics for 3,284 proteins were downloaded from the study's website. Cis-pQTL was searched within 500 kb distance from a target gene. Significant cis-eQTL was declared at p<5×10^−5^.

DEPICT is an integrative tool that takes advantage of predicted gene functions to systematically prioritize the most likely causal genes at loci of interest [27]. The input of DEPICT includes a list of variant identifiers, and the output contains all genes located in the loci and their p-values of being a candidate gene. All lead variants were submitted to DEPICT for analysis. Significant genes were declared at a false discovery rate (FDR)<5%.

SMR (Summary data–based Mendelian Randomization) method [61] is another SNP prioritization program that integrates summary-level data from GWAS with data from eQTL studies to identify genes whose expression levels are associated with trait due to causal or pleiotropy effects. Here, the pleiotropy effect means that a SNP is causally associated with both gene expression and phenotypic variation. SMR uses SNPs as an instrumental variable and tests the causal relation of gene expression to phenotype variation. The results are interpreted as the effect of gene expression on phenotype free of confounding from non-genetic factors. We used a pre-compiled eQTL dataset in whole blood tissue [62] for estimation.

### Gene-based and gene set enrichment analyses

Gene-based association analysis was performed with MAGMA v1.6 [63], as implemented on the FUMA website (http://fuma.ctglab.nl/). GWAS meta-analysis summary statistics were mapped to 19,427 protein-coding genes, resulting in 19,098 genes that were covered by at least one SNP. Gene-based association test was performed taking into account the LD between variants. Gene-based significance level was set at stringent Bonferroni corrected threshold 2.62×10^−6^, i.e., 0.05/19,098.

The generated gene-based summary statistics were further used to test for enrichment of association to specific biological pathways or gene sets. A gene set’s association signal was evaluated by integrating all signals from the genes in the set with MAGMA [63]. A competitive gene set analysis model was used to test whether the genes in a gene set are more strongly associated with the phenotype than other genes.

Gene sets were obtained through the MSigDB website (http://software.broadinstitute.org/gsea/msigdb/index.jsp) [64]. Each gene was assigned to a gene set as annotated by gene ontology (GO), Kyoto encyclopedia of genes and genomes (KEGG), Reactome and BioCarta gene set databases and other gene sets curated by domain experts or biomedical literature [64]. A total of 10,651 (4,734 curated and 5,917 GO terms) gene sets were used in this analysis. The significance level was set at a Bonferroni-corrected level of 0.05/10,651= 4.69×10^−6^.

Protein-protein interaction network was constructed with STRING [65]. STRING uses information based on gene co-expression, text-mining, and others, to construct protein interactive network.

### Polygenic risk score profiling

To assess the capability of the GWAS finding to predict ALM, a polygenic scoring analysis was conducted in the 369,968 unrelated subjects extracted from the main UKB sample. Three quarters of the individuals (277,762 subjects, including 149,329 females) were randomly selected as the training sample, and the remaining one quarter individuals (92,206 participants, including 49,660 females) as the validation sample. Female and male subjects were pooled together for analysis.

Raw phenotype was adjusted by age, age^2^, gender, height, height^2^ and the top 10 PCs, and the residuals were converted to the standard normal distribution quantiles for downstream analysis. Genetic association analysis was performed with PLINK2 [59].

The same QC procedures as in the main analysis were used to process the variants. The variants achieving a p-value of <1×10^−5^ in the training sample were selected and used for prediction in the validation sample via LDpred approach [66]. LDpred infers the posterior mean effect size of each marker by using a prior on effect sizes and LD information from an external reference panel. Specifically, the validation sample with original genotypes was used as reference panel for LD estimation. The number of SNPs used to adjust LD from each side of the target SNP was set to 1000. Other software parameters were set to the default.

### Genetic correlations with other traits

To test whether lean mass has a shared genetic etiology with other diseases and relevant traits, a genetic correlation analysis was performed with LDSC method [17]. An online web tool, LDHub (http://ldsc.broadinstitute.org/ldhub/), was used to estimate the genetic correlation between ALM_adj_ and 49 complex traits and diseases. The standalone version of the software was used to estimate between ALM_adj_ and two additional traits, ALM and total body lean mass, measured by the DXA scan, which are not available in the LDHub GWAS summary statistics collections, and which were downloaded from the GEFOS consortium website (http://www.gefos.org).

Both the LDHub and standalone analyses adopted same QC criteria. Specifically, only HapMap3 autosomal SNPs were included to minimize poor imputation quality [17]. SNPs were further removed given the following conditions: MAF<0.01, ambiguous strand (A/T or C/G), duplicated identifier, or reported sample size less than 60% of total sample size. LD scores pre-computed on the 1000 genomes project European individuals were used for calculation.

### Mendelian randomization analysis

To investigate whether ALM (as exposure) is causally associated with complex diseases (as outcome), a Mendelian randomization analysis with GSMR was performed [26] on selected 10 complex diseases, including fracture [36], type 2 diabetes (T2D) [37], asthma [38], insomnia [39], inflammatory bowel disease (IBD) [40], smoking addiction [41], coronary artery disease (CAD) [42], amyotrophic lateral sclerosis (ALS) [43], bipolar disorder [44] and autistic spectrum disorder (ASD) [45].

GWAS summary statistics for these diseases were downloaded from the respective websites. From the list of SNPs whose association signals with ALM_adj_ were below 5×10^−8^, qualified SNPs were included based on the following criteria: concordant alleles between exposure and outcome GWAS summary statistics, non-palindromic SNPs with certain strand, MAF>1%, and allele frequency difference between exposure and outcome GWAS summary statistics <0.2.

Independent SNPs were further clumped with PLINK2 [59] with independence LD threshold r^2^<0.05 and 1 MB window size. The clumped independent SNPs were examined for their pleiotropic effects to both exposure and outcome by the HEIDI test [26]. Significance level for the HEIDI test was set to α=1×10^−5^. After removing pleiotropic SNPs, the remaining independent SNPs were taken as instrumental variables to test for the causal effect of exposure to outcome. The estimated causal effect coefficients are approximately equal to the natural log odds ratio (OR) for a case–control trait. The MR analysis significance level was set to 0.005 (0.05/10).

## Acknowledgements

This research was conducted using the UK Biobank resource under application number 41542. This study was partially supported by the funding from national natural science foundation of China (31571291 to LZ, 31771417 and 31501026 to YFP), the natural science foundation of Jiangsu province of China (BK20150323 to YFP).

## References

1. Giles, J.T., et al., Association of body composition with disability in rheumatoid arthritis: impact of appendicular fat and lean tissue mass. Arthritis Rheum, 2008. 59(10): p. 1407–15.

2. Janssen, I., S.B. Heymsfield, and R. Ross, Low relative skeletal muscle mass (sarcopenia) in older persons is associated with functional impairment and physical disability. J Am Geriatr Soc, 2002. 50(5): p. 889–96.

3. Miyakoshi, N., et al., Prevalence of sarcopenia in Japanese women with osteopenia and osteoporosis. J Bone Miner Metab, 2013. 31(5): p. 556–61.

4. Binkley, N., D. Krueger, and B. Buehring, What’s in a name revisited: should osteoporosis and sarcopenia be considered components of “dysmobility syndrome?”. Osteoporos Int, 2013. 24(12): p. 2955–9.

5. Wannamethee, S.G. and J.L. Atkins, Muscle loss and obesity: the health implications of sarcopenia and sarcopenic obesity. Proc Nutr Soc, 2015. 74(4): p. 405–12.

6. Evans, W.J., et al., Cachexia: a new definition. Clin Nutr, 2008. 27(6): p. 793–9.

7. Brown, J.C., M.O. Harhay, and M.N. Harhay, Sarcopenia and mortality among a population-based sample of community-dwelling older adults. J Cachexia Sarcopenia Muscle, 2016. 7(3): p. 290–8.

8. Arden, N.K. and T.D. Spector, Genetic influences on muscle strength, lean body mass, and bone mineral density: a twin study. J Bone Miner Res, 1997. 12(12): p. 2076–81.

9. Livshits, G., et al., Contribution of Heritability and Epigenetic Factors to Skeletal Muscle Mass Variation in United Kingdom Twins. J Clin Endocrinol Metab, 2016. 101(6): p. 2450–9.

10. Liu, X.G., et al., Genome-wide association and replication studies identified TRHR as an important gene for lean body mass. Am J Hum Genet, 2009. 84(3): p. 418–23.

11. Ran, S., et al., Gene-based genome-wide association study identified 19p13.3 for lean body mass. Sci Rep, 2017. 7: p. 45025.

12. Hai, R., et al., Bivariate genome-wide association study suggests that the DARC gene influences lean body mass and age at menarche. Sci China Life Sci, 2012. 55(6): p. 516–20.

13. Urano, T., et al., Large-scale analysis reveals a functional single-nucleotide polymorphism in the 5’-flanking region of PRDM16 gene associated with lean body mass. Aging Cell, 2014. 13(4): p. 739–43.

14. Klimentidis, Y.C., et al., Genetic Variant in ACVR2B Is Associated with Lean Mass. Med Sci Sports Exerc, 2016. 48(7): p. 1270–5.

15. Zillikens, M.C., et al., Large meta-analysis of genome-wide association studies identifies five loci for lean body mass. Nat Commun, 2017. 8(1): p. 80.

16. Loh, P.R., et al., Efficient Bayesian mixed-model analysis increases association power in large cohorts. Nat Genet, 2015. 47(3): p. 284–90.

17. Bulik-Sullivan, B.K., et al., LD Score regression distinguishes confounding from polygenicity in genome-wide association studies. Nature Genetics, 2015. 47(3): p. 291–295.

18. Loh, P.R., et al., Contrasting genetic architectures of schizophrenia and other complex diseases using fast variance-components analysis. Nat Genet, 2015. 47(12): p. 1385–92.

19. Medina-Gomez, C., et al., Bivariate genome-wide association meta-analysis of pediatric musculoskeletal traits reveals pleiotropic effects at the SREBF1/TOM1L2 locus. Nat Commun, 2017. 8(1): p. 121.

20. Hubel, C., et al., Genomics of body fat percentage may contribute to sex bias in anorexia nervosa. Am J Med Genet B Neuropsychiatr Genet, 2018.

21. Finucane, H.K., et al., Partitioning heritability by functional annotation using genome-wide association summary statistics. Nat Genet, 2015. 47(11): p. 1228–35.

22. Finucane, H.K., et al., Heritability enrichment of specifically expressed genes identifies disease-relevant tissues and cell types. Nat Genet, 2018. 50(4): p. 621–629.

23. Consortium, G.T., Human genomics. The Genotype-Tissue Expression (GTEx) pilot analysis: multitissue gene regulation in humans. Science, 2015. 348(6235): p. 648–60.

24. Fehrmann, R.S., et al., Gene expression analysis identifies global gene dosage sensitivity in cancer. Nat Genet, 2015. 47(2): p. 115–25.

25. Sun, B.B., et al., Genomic atlas of the human plasma proteome. Nature, 2018. 558(7708): p. 73–79.

26. Zhu, Z., et al., Causal associations between risk factors and common diseases inferred from GWAS summary data. Nat Commun, 2018. 9(1): p. 224.

27. Pers, T.H., et al., Biological interpretation of genome-wide association studies using predicted gene functions. Nat Commun, 2015. 6: p. 5890.

28. Adzhubei, I.A., et al., A method and server for predicting damaging missense mutations. Nat Methods, 2010. 7(4): p. 248–9.

29. Kumar, P., S. Henikoff, and P.C. Ng, Predicting the effects of coding non-synonymous variants on protein function using the SIFT algorithm. Nat Protoc, 2009. 4(7): p. 1073–81.

30. Choi, Y., et al., Predicting the functional effect of amino acid substitutions and indels. PLoS One, 2012. 7(10): p. e46688.

31. Shihab, H.A., et al., Predicting the functional, molecular, and phenotypic consequences of amino acid substitutions using hidden Markov models. Hum Mutat, 2013. 34(1): p. 57–65.

32. Schmidt, K., et al., Sox8 is a specific marker for muscle satellite cells and inhibits myogenesis. J Biol Chem, 2003. 278(32): p. 29769–75.

33. Ge, X., et al., Lack of Smad3 signaling leads to impaired skeletal muscle regeneration. Am J Physiol Endocrinol Metab, 2012. 303(1): p. E90–102.

34. Estrella, N.L., et al., MEF2 transcription factors regulate distinct gene programs in mammalian skeletal muscle differentiation. J Biol Chem, 2015. 290(2): p. 1256–68.

35. Frank, N.Y., et al., Regulation of myogenic progenitor proliferation in human fetal skeletal muscle by BMP4 and its antagonist Gremlin. J Cell Biol, 2006. 175(1): p. 99–110.

36. Trajanoska, K., et al., Assessment of the genetic and clinical determinants of fracture risk: genome wide association and mendelian randomisation study. BMJ, 2018. 362: p. k3225.

37. Xue, A., et al., Genome-wide association analyses identify 143 risk variants and putative regulatory mechanisms for type 2 diabetes. Nat Commun, 2018. 9(1): p. 2941.

38. Moffatt, M.F., et al., A large-scale, consortium-based genomewide association study of asthma. N Engl J Med, 2010. 363(13): p. 1211–1221.

39. Hammerschlag, A.R., et al., Genome-wide association analysis of insomnia complaints identifies risk genes and genetic overlap with psychiatric and metabolic traits. Nat Genet, 2017. 49(11): p. 1584–1592.

40. Liu, J.Z., et al., Association analyses identify 38 susceptibility loci for inflammatory bowel disease and highlight shared genetic risk across populations. Nat Genet, 2015. 47(9): p. 979–986.

41. Genome-wide meta-analyses identify multiple loci associated with smoking behavior. Nat Genet, 2010. 42(5): p. 441–7.

42. van der Harst, P. and N. Verweij, Identification of 64 Novel Genetic Loci Provides an Expanded View on the Genetic Architecture of Coronary Artery Disease. Circ Res, 2018. 122(3): p. 433–443.

43. van Rheenen, W., et al., Genome-wide association analyses identify new risk variants and the genetic architecture of amyotrophic lateral sclerosis. Nat Genet, 2016. 48(9): p. 1043–8.

44. Stahl, E.A., et al., Genome-wide association study identifies 30 Loci Associated with Bipolar Disorder. bioRxiv, 2018: p. 173062.

45. Grove, J., et al., Identification of common genetic risk variants for autism spectrum disorder. Nat Genet, 2019. 51(3): p. 431–444.

46. Manolio, T.A., et al., Finding the missing heritability of complex diseases. Nature, 2009. 461(7265): p. 747–53.

47. Zuk, O., et al., The mystery of missing heritability: Genetic interactions create phantom heritability. Proc Natl Acad Sci U S A, 2012. 109(4): p. 1193–8.

48. Yang, J., et al., Common SNPs explain a large proportion of the heritability for human height. Nat Genet, 2010. 42(7): p. 565–9.

49. Wood, A.R., et al., Defining the role of common variation in the genomic and biological architecture of adult human height. Nat Genet, 2014. 46(11): p. 1173–86.

50. Consortium, S.W.G.o.t.P.G., Biological insights from 108 schizophrenia-associated genetic loci. Nature, 2014. 511(7510): p. 421–7.

51. Karasik, D. and D.P. Kiel, Genetics of the musculoskeletal system: a pleiotropic approach. J Bone Miner Res, 2008. 23(6): p. 788–802.

52. Blank, R.D., Bone and Muscle Pleiotropy: The Genetics of Associated Traits. Clin Rev Bone Miner Metab, 2014. 12(2): p. 61–65.

53. Loh, P.R., et al., Mixed-model association for biobank-scale datasets. Nat Genet, 2018. 50(7): p. 906–908.

54. Sanna, S., et al., Common variants in the GDF5-UQCC region are associated with variation in human height. Nat Genet, 2008. 40(2): p. 198–203.

55. Yang, J., et al., GCTA: a tool for genome-wide complex trait analysis. Am J Hum Genet, 2011. 88(1): p. 76–82.

56. Manichaikul, A., et al., Robust relationship inference in genome-wide association studies. Bioinformatics, 2010. 26(22): p. 2867–73.

57. Yengo, L., et al., Meta-analysis of genome-wide association studies for height and body mass index in approximately 700000 individuals of European ancestry. Hum Mol Genet, 2018. 27(20): p. 3641–3649.

58. Shungin, D., et al., New genetic loci link adipose and insulin biology to body fat distribution. Nature, 2015. 518(7538): p. 187–196.

59. Chang, C.C., et al., Second-generation PLINK: rising to the challenge of larger and richer datasets. Gigascience, 2015. 4: p. 7.

60. Benner, C., et al., Prospects of Fine-Mapping Trait-Associated Genomic Regions by Using Summary Statistics from Genome-wide Association Studies. Am J Hum Genet, 2017. 101(4): p. 539–551.

61. Zhu, Z., et al., Integration of summary data from GWAS and eQTL studies predicts complex trait gene targets. Nat Genet, 2016. 48(5): p. 481–7.

62. Westra, H.J., et al., Systematic identification of trans eQTLs as putative drivers of known disease associations. Nat Genet, 2013. 45(10): p. 1238–1243.

63. de Leeuw, C.A., et al., MAGMA: generalized gene-set analysis of GWAS data. PLoS Comput Biol, 2015. 11(4): p. e1004219.

64. Subramanian, A., et al., Gene set enrichment analysis: a knowledge-based approach for interpreting genome-wide expression profiles. Proc Natl Acad Sci U S A, 2005. 102(43): p. 15545–50.

65. Szklarczyk, D., et al., STRING v11: protein-protein association networks with increased coverage, supporting functional discovery in genome-wide experimental datasets. Nucleic Acids Res, 2019. 47(D1): p. D607–D613.

66. Vilhjalmsson, B.J., et al., Modeling Linkage Disequilibrium Increases Accuracy of Polygenic Risk Scores. Am J Hum Genet, 2015. 97(4): p. 576–92.

